# Persistent Inflammatory Lipotoxicity Impedes Pancreatic β-cell Function in Diet-Induced Obese Mice Despite Correction of Glucotoxicity

**DOI:** 10.1101/2022.05.31.494168

**Authors:** Ivan A. Valdez, Juan Pablo Palavicini, Terry M. Bakewell, Marcel Fourcaudot, Iriscilla Ayala, Ziying Xu, Ahmed Khattab, Xianlin Han, Chris E. Shannon, Luke Norton

## Abstract

Insulin resistance is a hallmark feature of Type 2 Diabetes (T2D), but the progression of the disease is closely linked to a deterioration in β-cell mass and function. While the precise mechanisms of β-cell failure are unclear, chronic hyperglycemia (glucotoxicity) and dyslipidemia (lipotoxicity) are considered contributing factors; however, the relative importance of these insults on β-cell function remains controversial. To examine this, we dissociated glucotoxicity from lipotoxicity using a high-fat diet (HFD)-fed mouse model of T2D and the glucose-lowering SGLT2 inhibitor, canagliflozin (CANA). As expected, HFD-feeding impaired glucose tolerance and isolated islet function. However, despite improvements in glucose tolerance and indices of β-cell insulin secretory function *in vivo*, CANA failed to restore isolated β-cell function. Shotgun lipidomics analysis of isolated islets revealed that HFD-feeding induced glycerophospholipid remodeling with a persistent increase in arachidonic acid (20:4)-enriched molecular species. Further analysis revealed that lysophosphatidylcholine (LPC) was the predominant lipid class elevated in HFD islets following correction of glucotoxicity with CANA. In follow-up experiments, LPC stimulations acutely and dose-dependently impaired glucose-stimulated insulin secretion (GSIS) in isolated wild-type islets, mechanistically linking this lipid class to β-cell dysfunction. Our findings indicate that persistent inflammatory lipotoxicity impedes β-cell function in diet-induced obese (DIO) rodents even after normalization of hyperglycemia. If replicated in humans, these data suggest that interventions targeting lipotoxicity may be beneficial for the long-term protection of pancreatic β-cell function in T2D.

## INTRODUCTION

Type 2 Diabetes (T2D) is characterized by hepatic and peripheral insulin resistance, concomitant β-cell hypersecretion, and a progressive deterioration in β-cell mass and function [1]. The increase in insulin secretion in the early stages of T2D is a compensatory response to excessive nutrient stress and may precede the detection of insulin resistance [2, 3]. Although the precise mechanisms linking nutrient excess to β-cell dysfunction in T2D are controversial, the direct toxic effects of high glucose (glucotoxicity) and/or lipids (lipotoxicity) on β-cell function are thought to be important contributors [4, 5].

Studies in rodents and human patients with T2D highlight a strong correlation between chronic hyperglycemia and the loss of β-cell mass and function. For example, the progression from impaired glucose tolerance (IGT) to T2D is associated with increased basal insulin, diminished first-phase glucose-stimulated insulin secretion (GSIS), and a gradual depletion of insulin stores [6-8]. Hyperglycemia-induced apoptosis is considered one of the major mechanisms promoting β-cell loss in T2D patients [9, 10], but β-cell dedifferentiation through the loss of insulin gene expression and reactivation of progenitor markers (e.g., Ngn3, Oct4, Nanog, and L-myc) may precede β-cell death [11-13]. Although it is unclear whether the detrimental effects of glucotoxicity on these pathways of β-cell loss can be fully reversed, therapies that alleviate hyperglycemia, promote weight loss, and/or increase insulin sensitivity (e.g., GLP-1 receptor agonists [14], sodium-glucose co-transporter 2 inhibitors (SGLT2i) [15], thiazolidinediones (TZDs) [16], and insulin therapy [17]) have been shown to enhance markers of beta cell identity or reverse markers of β-cell dedifferentiation.

There is less consensus on whether lipotoxicity is an independent driver of β-cell dysfunction in T2D. Much of the supportive evidence is derived from *in vitro* islet studies, which have linked supra-physiological doses of specific fatty acids to defects in β-cell function [18-20]. While the accurate quantitation of physiological lipid species and concentrations within or surrounding islets in various disease states *in vivo* remains a challenge [5], several published findings support a role for lipotoxicity in β-cell dysfunction in animals and humans. For example, temporal diet-induced increases in circulating free fatty acid (FFA) levels in rodent models of T2D are closely correlated with the onset of β-cell dysfunction [21]. FFAs proximal to β-cells are important modulators of GSIS, and increased hydrolysis of triacylglycerols (TAGs) and delivery of FFAs to β-cells via lipoprotein lipase (LPL) overexpression in the pancreas impairs β-cell function and promotes glucose intolerance [22]. In humans, plasma TAGs are closely correlated with the expression of genes controlling β-cell mass and function [23], and chronic FFA infusion impairs the acute stimulatory effects of FFAs on insulin secretion [24]. Finally, ectopic lipid deposition in the whole pancreas has been observed in patients with obesity and T2D using ^1^H magnetic resonance spectroscopy [25, 26], although the causal link between pancreatic steatosis and β-cell function has not been firmly established and remains controversial [27, 28].

Dissecting the relative contributions of gluco-, lipo- and glucolipo-toxicities to β-cell dysfunction is a clinically significant area of investigation because maintaining long-term β-cell health is crucial for the prevention and treatment of T2D. In the present study, we sought to examine the relative contribution of glucotoxicity and lipotoxicity *in vivo* by treating diet-induced obese (DIO) mice with canagliflozin (CANA), a highly specific inhibitor of sodium-glucose co-transporter 2 (SGLT2) in the kidney. The SGLT2 inhibitor (SGLT2i) class of medications alleviates glucotoxicity by blocking the reuptake of filtered glucose through SGLT2 and promoting urinary glucose excretion. To quantitate lipotoxicity directly in β-cells, we developed a multi-dimensional mass spectrometry-based shotgun lipidomics technique in islets isolated from mice.

Our study shows that 5 weeks of CANA treatment improved body weight and alleviated hyperglycemia, as expected, but failed to restore isolated β-cell function in high-fat diet (HFD)-fed mice. Islet lipidomics revealed significant glycerophospholipid remodeling in HFD mouse islets, with a marked increase in pro-inflammatory arachidonic acid (20:4)-enriched molecular species. Lysophosphatidylcholine (LPC), a pro-inflammatory lipid clinically linked to insulin resistance and T2D [29], remained elevated in HFD islets despite correction of glucotoxicity with CANA. In follow-up experiments using wild-type islets, LPC acutely and dose-dependently increased basal insulin secretion and impaired GSIS, mechanistically linking this pro-inflammatory lipid to β-cell dysfunction. Together, these findings demonstrate that disturbances in the islet lipidome negatively impact β-cell function *in vivo* and *in vitro*, and they highlight the potential importance of targeting both hyperglycemia and dyslipidemia to preserve β-cell health in patients with T2D.

## RESULTS

### SGLT2i Abrogates Glucotoxicity in Diet-Induced Obese (DIO) Mice

For this study, male C57BL/6 mice were fed a regular chow (RC) or high-fat diet (HFD) for 8 weeks. After 8 weeks of diet, RC mice were treated with vehicle (cellulose; 10 mg/kg body weight/day), and mice fed the HFD were treated with either vehicle or the SGLT2i, canagliflozin (CANA; 10 mg/kg body weight/day), which was administered daily via oral gavage. Thus, the three groups used in this study were: 1) RC mice treated with vehicle (RC), 2) HFD fed mice treated with vehicle (HFD), and 3) HFD fed mice treated with canagliflozin (HFD CANA) (**Fig. 1A**).

**Figure 1.**
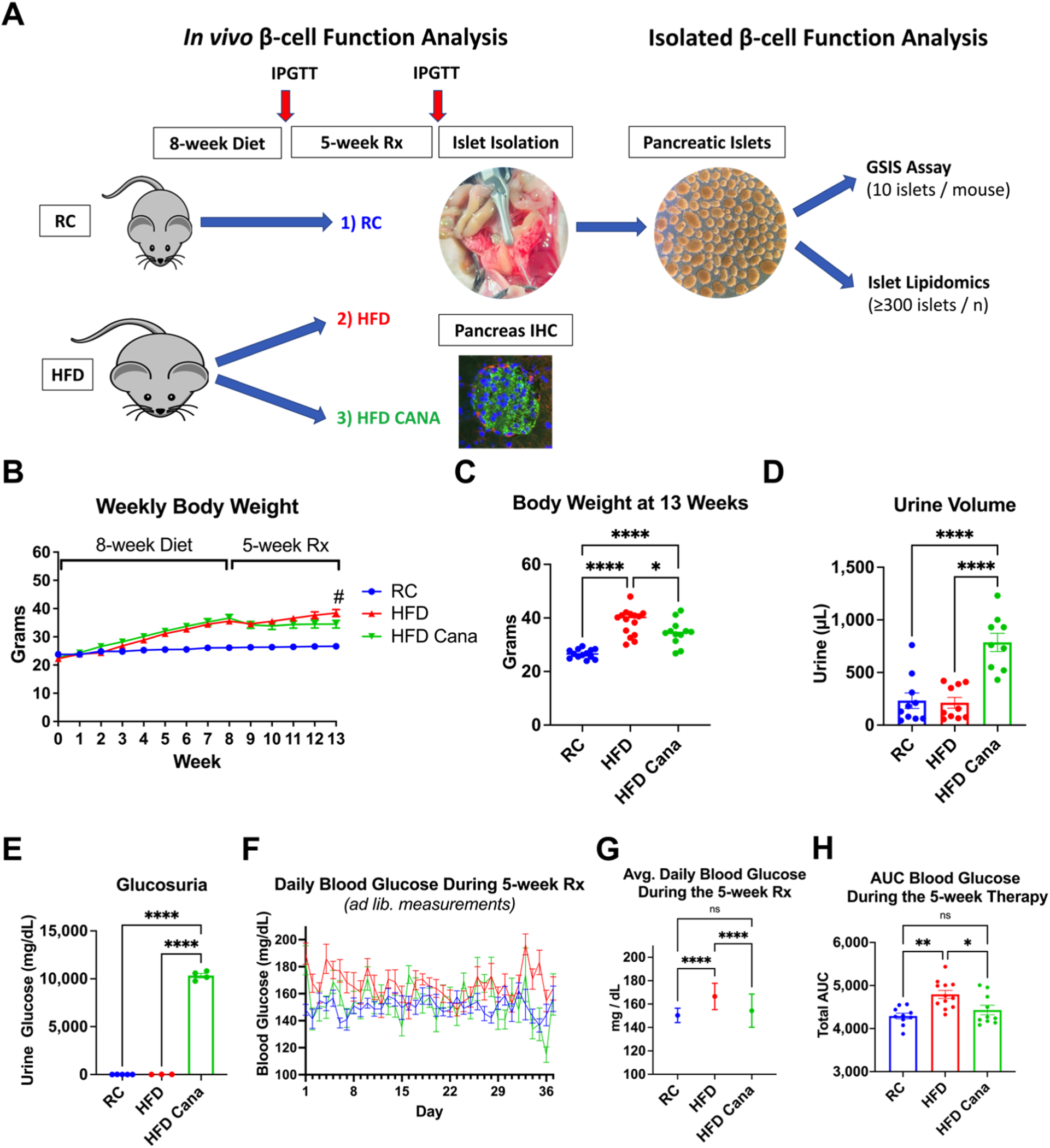
SGLT2i Abrogates Glucotoxicity in DIO Mice. **(A)** Experimental design: Male C57BL/6 mice were fed a regular chow (RC) or high-fat diet (HFD) for 8 weeks followed by 5 weeks of vehicle (VEH: cellulose, 10 mg/kg body weight/day) or the SGLT2i, canagliflozin (CANA; 10 mg/kg body weight/day) administered daily via oral gavage. Pancreases were then harvested for immunohistochemistry (IHC) or islet isolation via collagenase technique for glucose-stimulated insulin secretion (GSIS) or mass spectrometry-based shotgun lipidomics. **(B)** Weekly body weight measurements (n=12-15 per group). **(C)** Body weight at 13 weeks of age (n=12-15 per group). **(D)** Urine volumes collected 5 hours after VEH of CANA administration (n=9-10 per group). **(E)** Urine glucose levels (n=3-5 per group). **(F)** Daily ad libitum blood glucose levels during the 5-week treatment (n=10-12 per group). **(G)** Average daily blood glucose levels during the 5-week treatment (n=10-12 per group). **(H)** Total area under the curve (AUC) for daily blood glucose levels during the 5-week treatment **(Fig. 1F)** (n=10-12 per group). Data represent mean +/- SEM; ns (no significance), *p≤0.05, **p≤0.01, ***p≤0.005, ****p≤0.0005; HFD vs. HFD CANA: #p≤0.05. Data analyzed using an ordinary 1-way or 2-way ANOVA, where appropriate, with correction for multiple comparisons using Holm-Šídák’s method.

After 8-weeks of diet, HFD mice were approximately 10 g (∼40%) heavier than RC mice, on average (**Fig. 1B**). As expected, and consistent with data from human patients with T2D and prior animal studies [30, 31], HFD CANA mice lost approximately 10% (∼4g) of their bodyweight following 5 weeks of treatment but remained significantly heavier than RC mice at the end of the treatment period (**Fig. 1B** and **Fig. 1C**). To examine the efficacy of SGLT2i therapy on glycemic levels, we confirmed significant urinary glucose excretion in mice treated with CANA (**Fig. 1D** and **Fig. 1E**) and measured blood glucose daily during the treatment period. As shown in **Fig. 1F – 1H**, HFD feeding increased average daily blood glucose and glucose area under the curve (AUC), both of which were completely normalized in mice treated with CANA.

### SGLT2i Improves Glucose Tolerance and Restores Insulin Sensitivity

To quantitate longitudinal glucose homeostasis, we performed two rounds of intraperitoneal glucose tolerance tests (IPGTTs); before and after CANA treatment (**Fig. 1A**). We used a fixed dose of glucose (50 mg) for all IPGTTs irrespective of body weight. The first round of IPGTTs was performed after 8 weeks of RC or HFD feeding. As expected, HFD mice demonstrated increased fasting blood glucose levels and glucose intolerance during the IPGTT (**Fig. 2A** and **Fig. 2B**). The incremental AUC (iAUC) during the IPGTT was also significantly higher in HFD-fed mice, indicating the presence of insulin resistance in HFD-fed mice prior to CANA treatment (**Fig. 2C**). The second round of IPGTTs was performed 5 weeks after vehicle or CANA therapy and, consistent with the data described above, CANA normalized glucose tolerance (**Fig. 2D – 2F**).

**Figure 2.**
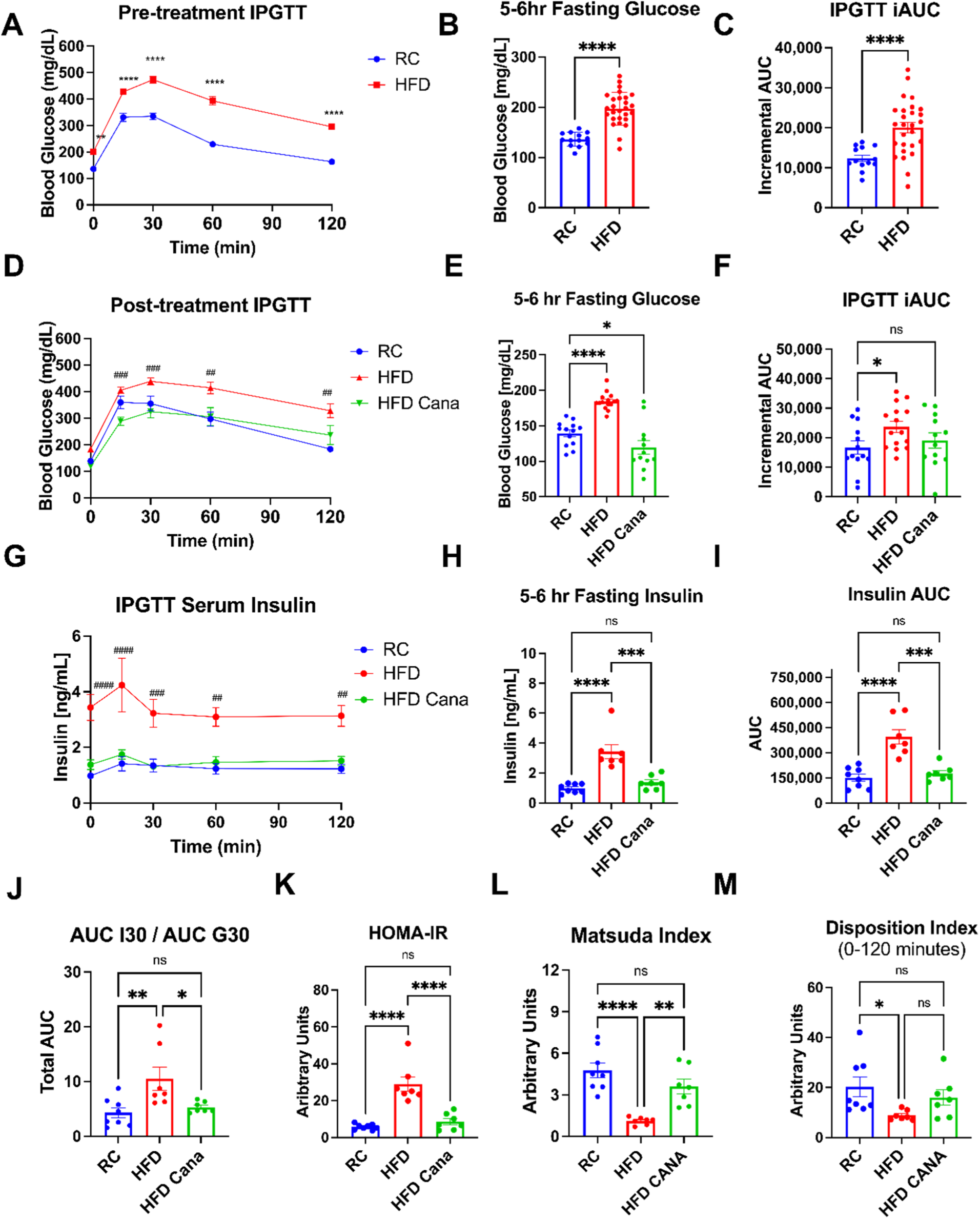
SGLT2i Improves Glucose Tolerance, Insulin Sensitivity, and *In Vivo* β-Cell Function. **(A)** Pre-treatment intraperitoneal glucose tolerance test (IPGTT) using a fixed dose of 50mg of glucose (500µL of 10% dextrose solution) after a 5-6 hour fast (RC, n=13; HFD, n=27). **(B)** Fasting (5-6hr) blood glucose levels during pre-treatment IPGTT (RC, n=13; HFD, n=27). **(C)** Incremental area under the curve (iAUC; normalizing for baseline differences) for pre-treatment IPGTT (RC, n=13; HFD, n=27) **(D)** Post-treatment IPGTT using a fixed dose of 50mg of glucose (500µL of 10% dextrose solution) after a 5-6 hour fast (n=12-15 per group). **(E)** Fasting (5-6hr) blood glucose levels during the post-treatment IPGTT (n=12-15 per group). **(F)** iAUC for blood glucose levels during the post-treatment IPGTT (n=12-15 per group). **(G)** Serum insulin levels during the post-treatment IPGTT (n=7-8 per group). **(H)** Fasting (5-6hr) serum insulin levels during the post-treatment IPGTT (n=7-8 per group). **(I)** Total AUC for serum insulin during the post-treatment IPGTT (n=7-8 per group). **(J)** Ratio of the 30-minute AUC for insulin (AUC I30; **Supp. Fig. 1A**) factored by the 30-minute AUC for glucose (AUC G30; **Supp. Fig. 1B**); (n=7-8 per group). **(K)** HOMA IR measurement: (Fasting Insulin) x (Fasting Glucose) / (22.5 × 1,000); (n=7-8 per group). **(L)** Matsuda Index: (10^6/square root of [fasting glucose x fasting insulin] x [mean glucose x mean insulin during OGTT]) as described by Matsuda and DeFronzo [94] (n=7-8 per group). **(M)** Disposition Index during the post-treatment IPGTT: (AUC Insulin 0-120 / AUC Glucose 0-120) x (Matsuda Index); (n=7-8 per group). Data represent mean +/- SEM, ns (no significance), *p≤0.05, **p≤0.01, ***p≤0.001, ****p≤0.0001; ##p≤0.01, ###p≤0.001, ####p≤0.0001 HFD vs HFD CANA; Data analyzed using an unpaired t-test with Welch’s correction and an ordinary 1-way or 2-way ANOVA, where appropriate, with correction for multiple comparisons using Holm-Šídák’s method.

### SGLT2i Improves *In Vivo* β-Cell Function in DIO Mice

Human and rodent studies have demonstrated improvements in β-cell function following treatment with SGLT2i *in vivo*, which has been linked mechanistically to improved glucotoxicity [32-37]. However, it is not clear whether this reflects protection of individual β-cells against progressive β-cell decline in T2D or a compensatory response to the favorable metabolic milieu *in vivo* following SGLT2i therapy. To determine this, we examined indices of β-cell function *in vivo* and quantitated β-cell function directly in islets isolated from mice treated with CANA.

To assess *in vivo* insulin secretion, we collected serum during the post-treatment IPGTT and measured insulin levels. Mice fed the HFD exhibited significant hyperinsulinemia across all timepoints of the IPGTT compared to RC mice, which was completely normalized in HFD CANA mice (**Fig. 2G – 2I**). The 30-minute AUC for insulin (AUC I30; **Supp. Fig. 1A**) factored by the 30-minute AUC for glucose (AUC G30; **Supp. Fig. 1B**), which quantifies the initial phase of insulin secretion, revealed a disproportionate increase in insulin secretion in HFD mice that was corrected in the HFD CANA group (**Fig. 2J**). Because *in vivo* β-cell function is significantly influenced by insulin resistance, we used HOMA-IR (**Fig. 2K**) and the Matsuda Index (**Fig. 2L**) to demonstrate that insulin sensitivity was markedly improved in HFD CANA mice. We then estimated β-cell function using the Disposition Index, which evaluates β-cell function relative to insulin sensitivity [38]. Using this measure, β-cell function was clearly impaired in mice fed the HFD, but was improved by approximately 80% following CANA treatment, although this did not reach statistical significance (p = 0.2; **Fig. 2M**). These data demonstrate that correction of glucotoxicity using SGLT2i markedly improves glucose tolerance, decreases insulin resistance, and largely restores β-cell insulin secretory function *in vivo*.

Immunohistochemistry (IHC) analysis revealed the presence of SGLT1 protein in pancreatic sections of our experimental groups (**Supp. Fig. 2A)** and isolated wild-type islets (**Supp. Fig. 2B**), as has been previously reported in rodent [39, 40] and human [41] pancreatic α-cells. SGLT2 protein was not detected, confirming recent reports [41, 42].

### SGLT2i Does Not Rescue Isolated β-Cell Function

To examine the function of isolated β-cells, we carried out an intraductal collagenase perfusion procedure to isolate pancreatic islets and performed glucose-stimulated insulin secretion (GSIS) assays, the gold-standard method used to measure β-cell function (**Fig. 3A**). We first quantified islet size distribution across the three experimental groups. HFD-feeding led to a marked increase in average islet size, which was largely reversed by CANA treatment (**Fig. 3B – 3E**).

**Figure 3.**
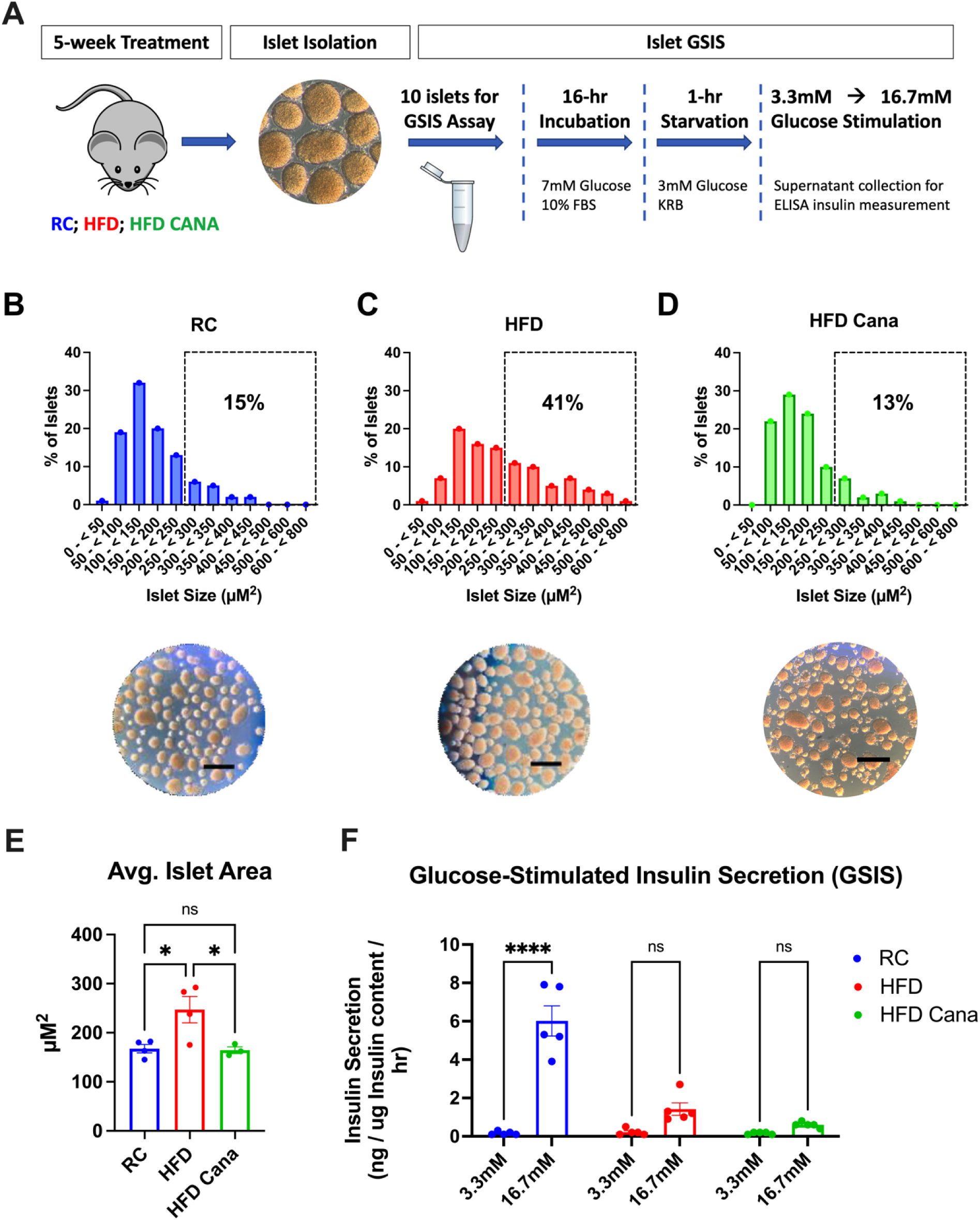
SGLT2i Does Not Rescue Isolated β-Cell Function. **(A)** Schematic of glucose-stimulated insulin secretion (GSIS) experimental design. **(B-D)** Size distribution and representative images of isolated islets (n=3-4 per group). Scale = 400µM. **(E)** Average islet size of isolated islets (n=3-4 per group). **(F)** Insulin levels during GSIS using low [3.3mM] or high [16.7mM] glucose (n=5 per group). GSIS data confirmed with 2 independent experiments using independent mouse groups/treatments (**Supp. Fig. 3C**). Data represent mean +/- SEM, ns (no significance), *p≤0.05, **p≤0.01, ***p≤0.005, ****p≤0.0005. Data analyzed using an ordinary 1-way or 2-way ANOVA, where appropriate, with correction for multiple comparisons using Bonferroni’s method.

We then hand-picked size-matched (**Supp. Fig. 3A**) islets for quantitation of GSIS *in vitro*. As expected, GSIS was significantly impaired in islets isolated from HFD mice (**Fig. 3F**). Remarkably, however, and in contrast to our *in vivo* data, SGLT2i treatment failed to restore glucose sensitivity in isolated islets (**Fig. 3F**), despite confirmation that depolarization-driven insulin secretion with 30 mM KCL remained intact and consistent across all three groups (**Supp. Fig. 3B**). To confirm these findings, we repeated our experiments in an independent cohort of mice fed a HFD and treated with VEH or CANA. Consistent with our initial experiments in **Fig. 3F**, the impairment in GSIS in islets isolated from this cohort of HFD-fed mice also was refractory to CANA therapy (**Supp. Fig. 3C**). It is clear from these data that, while β-cell glucose sensitivity is responsive to SGLT2i treatment *in vivo*, the function of isolated islets remains significantly impaired following the correction of glucotoxicity.

### SGLT2i Decreases Pancreatic CD68+ Macrophage Infiltration

In addition to glucotoxicity, localized islet inflammation has been associated with human and rodent β-cell failure through the presence of pancreatic macrophage infiltration, amyloid deposits, and fibrosis [43-46]. To assess macrophage-mediated inflammation as a potential mechanism of β-cell dysfunction, we performed immunohistochemistry (IHC) analyses by co-staining macrophage-specific marker, CD68, with insulin (**Fig. 4A**). Indeed, pancreatic islets from HFD mice revealed a significant increase in localized CD68+ cells (**Fig. 4B**), the percentage of islets with 5 or more CD68+ cells (**Fig. 4C**), and the number of CD68+ cells in acinar tissue surrounding islets (**Fig. 4D**). However, these effects were all restored by CANA treatment (**Fig. 4B – 4D**). Secondary antibody negative controls were used to rule out non-specific staining (**Supp. Fig. 4A**).

**Figure 4.**
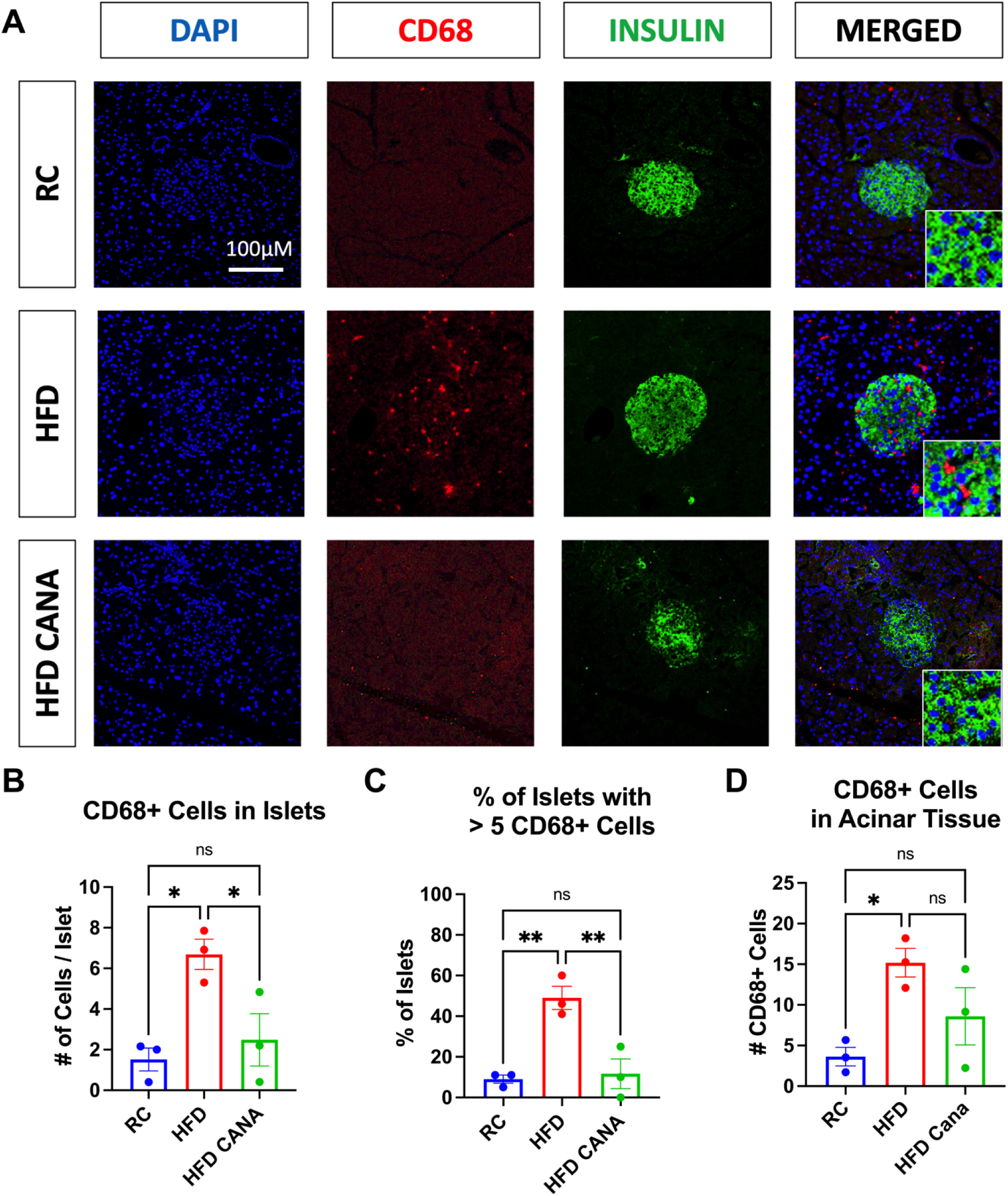
SGLT2i Decreased Pancreatic CD68+ Macrophage Infiltration. **(A)** Representative images of immunohistochemistry (IHC) experiments on pancreatic sections; macrophage marker, CD68 (Red), Insulin (green); DAPI (blue). Scale = 100μM. **(B)** Average number of CD68+ cells per islet. **(C)** Percentage of islets that contain 5 or more CD68+ cells. **(D)** Average number of CD68+ cells in acinar tissue surrounding pancreatic islets observed in 20X resolution. Data represent mean +/- SEM, ns (no significance), *p≤0.05, **p≤0.01, ***p≤0.005, ****p≤0.0005 (n=4-6 per group; a minimum of 20 islets were analyzed per mouse). Data analyzed using an ordinary 1-way ANOVA with correction for multiple comparisons using Holm-Šídák’s method.

### Islet Lipidomics Identified >200 Molecular Species Across 16 Functional Lipid Classes

To examine alternative mechanisms that could underlie isolated β-cell dysfunction following correction of glucotoxicity, we developed a methodology to profile the lipidome in a small number of islets isolated from mice. Preliminary lipidomics optimization experiments using C57BL/6 wild-type mouse islets showed that a minimum of 250-300 islets were needed to successfully detect lipid species of interest. The number of islets (∼300-400 per sample) we used for each of our three experimental groups is listed in **Supp. Table 1**.

Our class-targeted shotgun lipidomics approach determined the islet lipid content of 240 molecular lipid species (**Supp. Fig. 5A – 5D**) across 16 functional lipid classes (**Fig. 5A – 5B)**. These 16 lipid classes can be further divided into 4 structural families: 1) Sphingolipids (**Fig. 5C**) 2) Fatty Acyl Lipids (**Fig. 5D**); 3) Glycerolipids (**Fig. 5E**); and 4) Glycerophospholipids (**Fig. 5F**).

**Figure 5.**
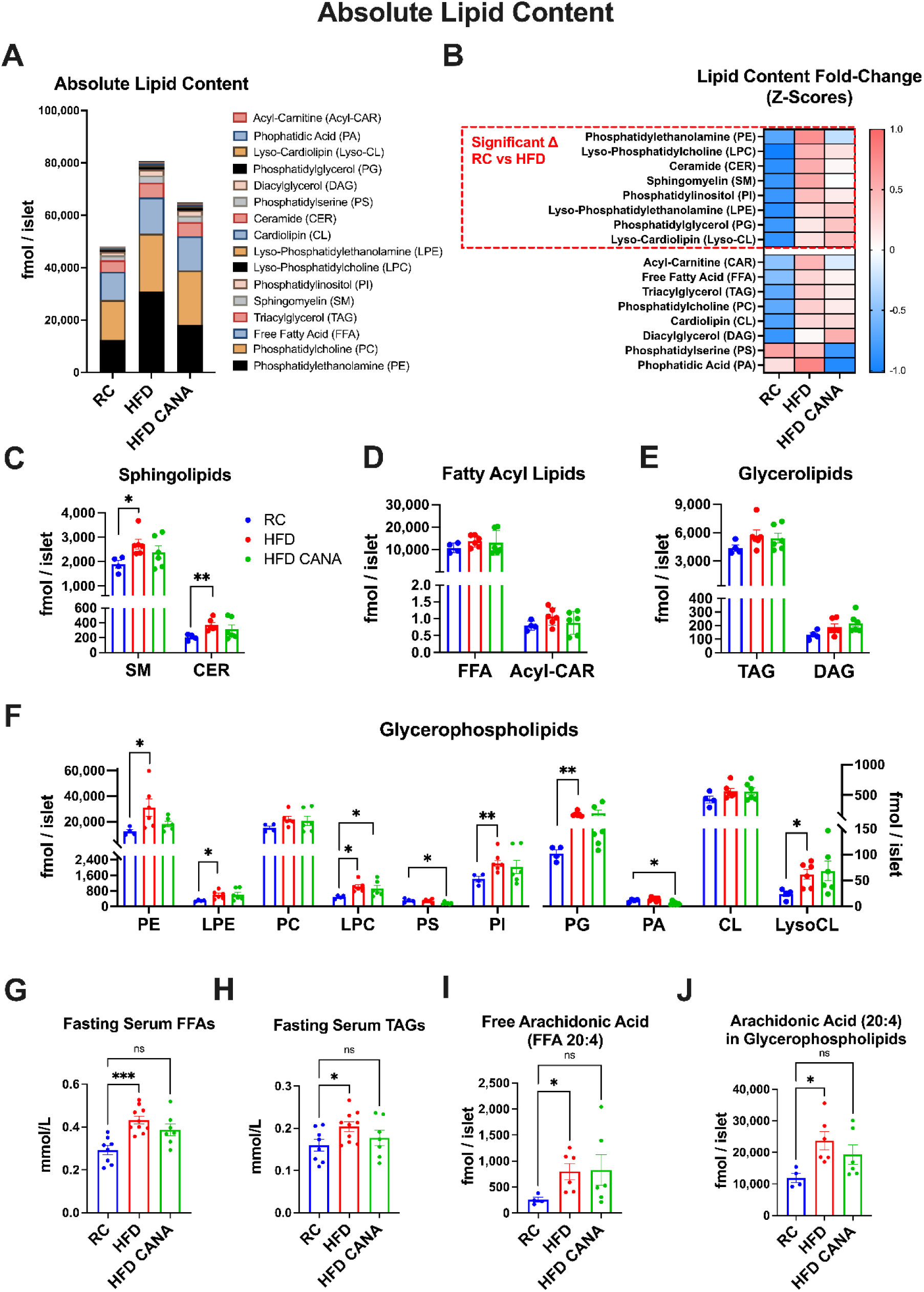
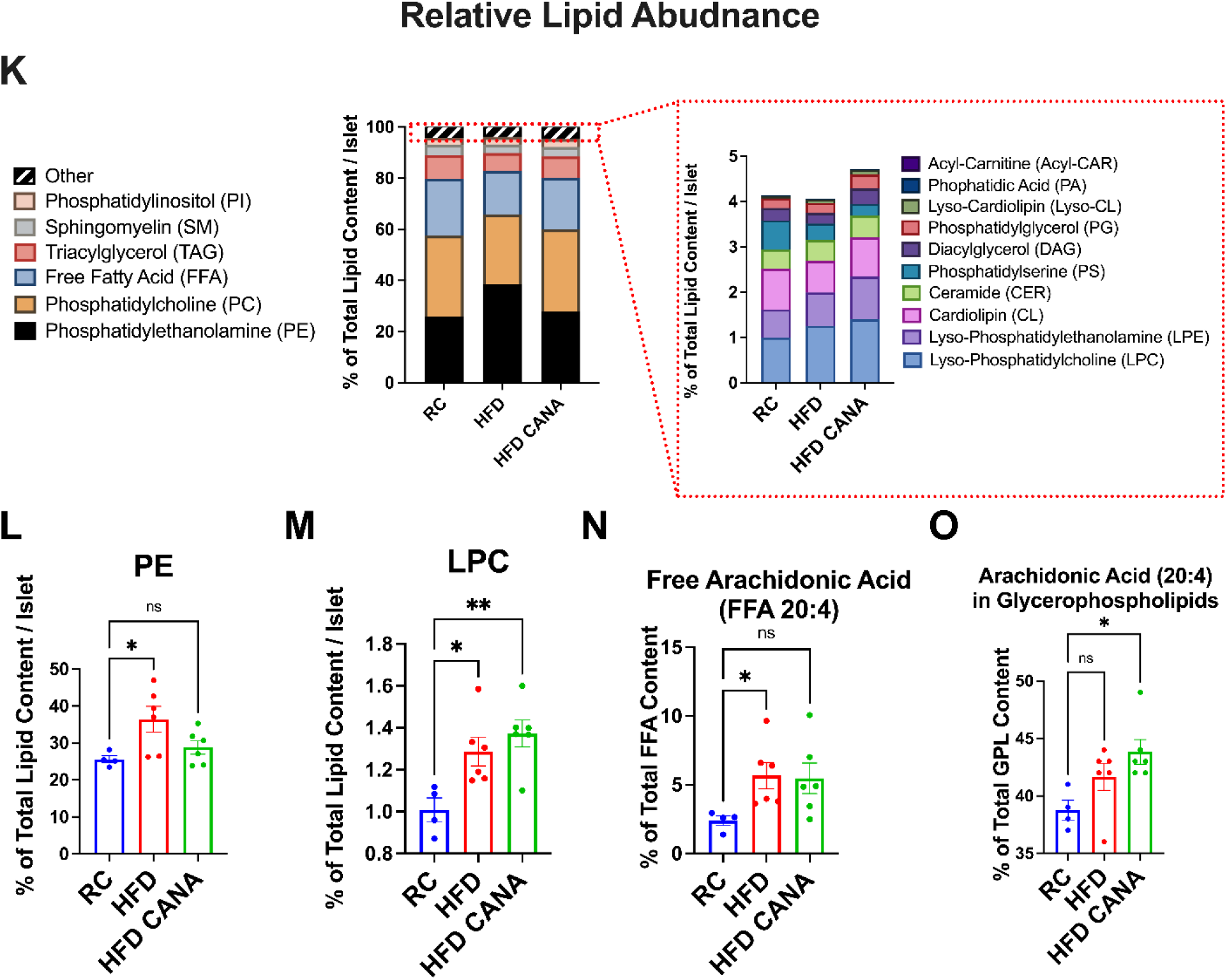
Lipidomics Reveals Persistent Inflammatory Lipotoxicity in HFD CANA Islets. **(A)** Absolute lipid content for 16 lipid classes measured expressed in fmol/islet. **(B)** Heatmap depicting fold change (z-scores) for all 16 molecular lipid classes in isolated islets from RC, HFD, or HFD CANA mice. Significantly altered lipids (p < 0.05 using Brown-Forsythe 1-way ANOVA, with correction for multiple comparisons using Dunnett’s method) inside dotted box, sorted by descending fold change between HFD vs RC. **(C)** Absolute lipid content of Sphingolipids: sphingomyelin (SM), ceramide (CER). **(D)** Absolute lipid content of Fatty Acyl Lipids: free fatty acids (FFA), acylcarnitine (Acyl-CAR). **(E)** Absolute lipid content of Glycerolipids: triglycerides (TAG), diacylglyceride (DAG). **(F)** Absolute lipid content of Glycerophospholipids and Lyso-glycerophospholipid intermediates: phosphatidylethanolamine (PE), lyso-phosphatidylethanolamine (LPE), phosphatidylcholine (PC), lyso-phosphatidylcholine (LPC), phosphatidylserine (PS), phosphatidylinositol (PI), phosphatidylglycerol (PG), phosphatidic acid (PA), cardiolipin (CL), and lyso-cardiolipin (Lyso-CL). **(G)** 12-hour fasting serum free fatty acids (FFAs). **(H)** 12-hour fasting serum triacylglycerol/triglycerides (TAGs). **(I)** Absolute content of free arachidonic acid (FFA 20:4) expressed as fmol/islet. **(J)** Absolute content of arachidonic acid (20:4) enrichment in glycerophospholipids expressed as fmol/islet. **(K)** Relative abundance of the 16 lipid classes as a percentage of total lipid content per islet. **(L)** Relative abundance of PE. **(M)** Relative abundance of LPC. **(N)** Relative abundance of free arachidonic acid (FFA 20:4) expressed as percentage of total FFA content/islet. **(O)** Relative abundance of arachidonic acid (20:4) enrichment in glycerophospholipids expressed as percentage of total GPL (glycerophospholipid) content/islet. Data represent mean +/- SEM, ns (no significance), *p≤0.05, **p≤0.01, ***p≤0.005, ****p≤0.0005; (n=4-6 per group for all figures); Data analyzed using Brown-Forsythe 1-way ANOVA, with correction for multiple comparisons using Dunnett’s method. All data normalized to islet number.

### HFD-feeding Induced Islet Glycerophospholipid Remodeling

We first examined changes in islet lipids in response to HFD-feeding. Although the increase in total lipid content measured in HFD islets did not attain statistical significance (p = 0.07; **Fig. 5A**), 8 of 16 lipid classes identified were significantly increased (**Fig. 5B – 5F**).

Several of the lipids increased in HFD islets are linked to inflammatory processes. For example, sphingomyelin and ceramides (**Fig. 5C**) have both been associated with β-cell lipotoxicity and insulin secretory defects in isolated rodent islets, as well as obesity, insulin resistance, and T2D in humans [47-49]. Most HFD-induced changes occurred in glycerophospholipids (**Fig. 5F**), which are critical components of cell membrane fluidity and stability [50]. Phosphatidylethanolamine (PE) was the most abundant lipid measured (**Fig. 5A**), and it showed the greatest fold-increase in HFD islets (**Fig. 5B**). It is also known to contain the largest portion of plasmalogens—molecular species with an ether linkage, rather than an acyl linkage, at the sn-1 position that are highly enriched in arachidonic acid (20:4) [51]. In states of metabolic stress, arachidonic-acid (20:4)-containing plasmalogens become precursors of inflammatory mediators (e.g., leukotrienes and hydroxy-eicosatetraenoic acids) [50], and therefore contribute to the inflammatory response. Indeed, our lipidomics analysis demonstrated that the increase in total PE was driven by plasmalogen species (P-PE) rather than diacyl species (D-PE) (**Supp. Fig. 5D**). Other glycerophospholipids and lysoglycerophospholipid intermediates increased by HFD include: lyso-phosphatidylcholine (LPC), lyso-phosphatidylethanolamine (LPE), lyso-cardiolipin (LysoCL), phosphatidylglycerol (PG), and phosphatidylinositol (PI) (**Fig. 5B and 5F**). These findings are noteworthy since alterations in glycerophospholipid composition has been reported in several metabolic and inflammatory diseases [52, 53].

Islet FFAs (**Fig. 5D**) and glycerolipids (i.e., TAGs and DAGs) (**Fig. 5E**) were surprisingly not increased in HFD islets, despite increased levels of fasting serum FFAs (**Fig. 5G**) and TAGs (**Fig. 5H**) in HFD mice. Furthermore, even though TAGs and DAGs make up a significant portion of total lipids in other tissues (e.g., adipose tissue [54] or liver [55]), these lipids were generally not as abundant in pancreatic islets (**Fig. 5A**).

Lastly, we quantified the relative abundance of each lipid class as a percentage of total lipid measured (**Fig. 5K**). This analysis showed that PE and LPC were significantly increased in HFD islets relative to other lipids (**Fig. 5L and 5M**; **Supp. Fig. 6A – 6D**). Phosphatidylserine (PS), an inner phospholipid that is externalized in states of metabolic stress and apoptosis [56], and phosphatidic acid (PA), the obligate precursor for all glycerophospholipids [54], tended to decrease in HFD-feeding conditions in terms of absolute (**Fig. 5F**) and relative abundance (**Supp. Fig. 6D**). Together, these findings suggest an association between HFD-feeding and islet glycerophospholipid remodeling.

### HFD-feeding Increased Arachidonic Acid (20:4)-enriched Molecular Species

Many of the individual molecular species that increased in HFD-fed mouse islets contained arachidonic acid (20:4), and most belonged to the glycerophospholipid family (**Supp. Fig 5A – 5F**). Our analysis confirmed that free arachidonic acid was increased in HFD islets in terms of absolute content (**Fig. 5I**) and relative abundance (**Fig. 5N**). Arachidonic acid-enrichment in glycerophospholipids was also significantly increased in terms of absolute content (**Fig. 5J**) and showed a strong, but non-significant increase in relative abundance (**Fig. 5O**). Since arachidonic acid serves as a precursor to various signaling lipids related to inflammation and has been associated with beta cell dysfunction [57, 58], these lipidomics findings suggest that inflammatory lipotoxicity is a potential mechanism underlying HFD-mediated β-cell dysfunction.

### Lipidomics Reveals Persistent Inflammatory Lipotoxicity in HFD CANA Islets

We next analyzed the effects of CANA on islet lipids. To identify lipid classes and molecular species that contribute to persistent β-cell dysfunction in DIO mice following glycemic control, we divided lipids into those that responded to CANA treatment and those that did not.

Free arachidonic acid (20:4) was not significantly elevated in HFD CANA islets relative to RC in terms of absolute content (p = 0.2; **Fig. 5I**) or relative abundance (p = 0.07; **Fig. 5N**), despite an upward trend. Arachidonic acid-enrichment in glycerophospholipid species also trended upwards in HFD islets in terms of absolute content (**Fig. 5J**) and was significantly increased in terms of relative abundance (**Fig. 5O**). These data suggest that arachidonic acid was largely normalized in HFD CANA islets but may contribute to β-cell dysfunction as a component of distinct glycerophospholipid species.

Out of the 8 lipid classes that increased after HFD-feeding, LPC remained elevated in HFD CANA islets in terms of absolute content (**Fig. 5F**) and relative abundance (**Fig. 5M**). This increase was largely driven by 4 of 12 molecular species (16:0, 18:0; 18:1, and 20:4), which collectively represented ∼87% of total LPC content measured (**Fig. 6A**). Arachidonic acid (20:4)-enrichment in LPC remained significantly elevated in HFD CANA islets (**Fig. 6A**). Based on this data and, given that LPC levels are increased in the serum of patients with T2D [29, 59, 60], we sought to further explore the effects of LPC on isolated β-cell function.

**Figure 6.**
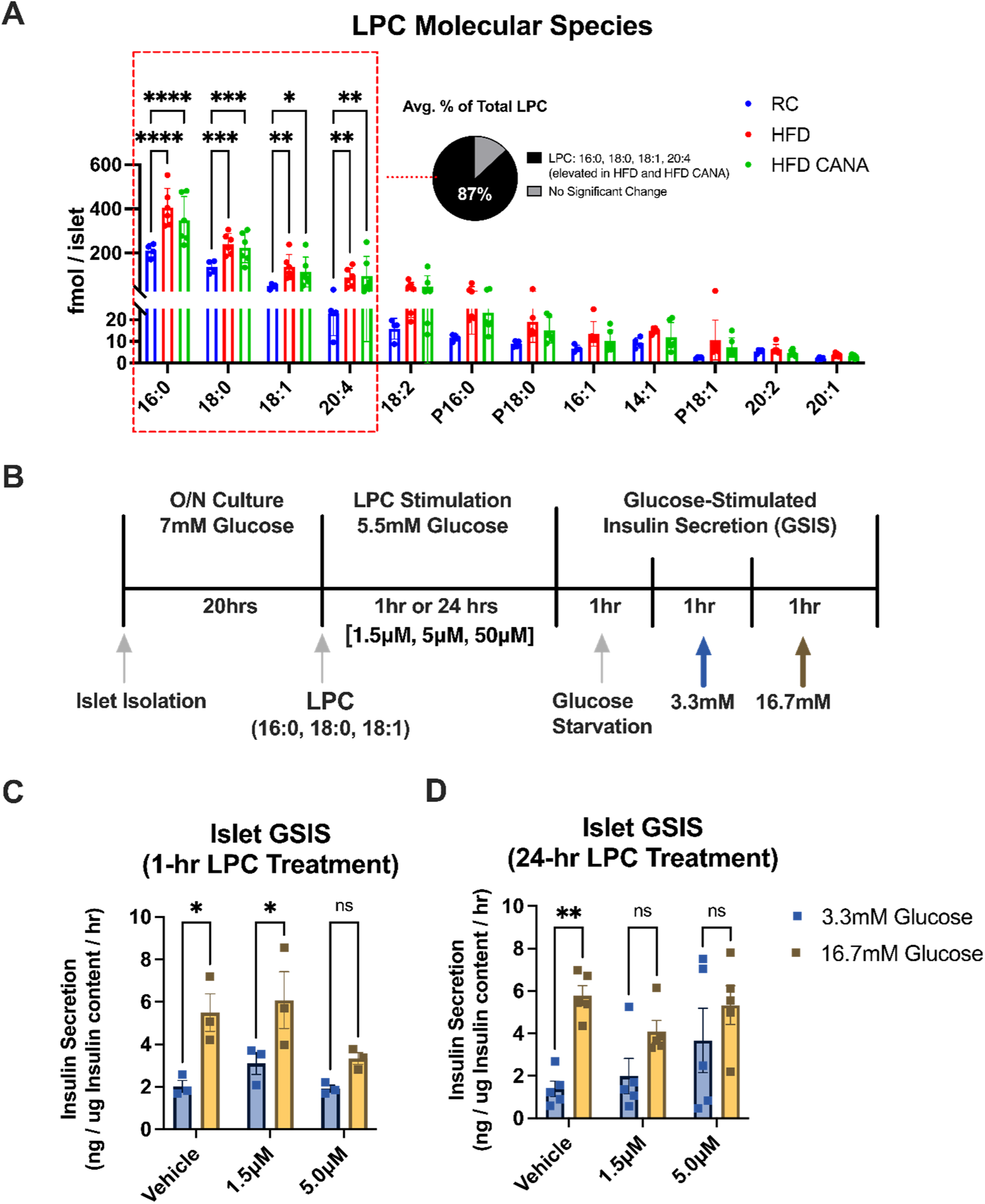
LPC Stimulation Disrupts β-cell Function Through Impaired GSIS. **(A)** LPC content by individual molecular species. Pie chart depicts average percentage of total LPC content of the 4 species that had significant differences in RC vs HFD and HFD CANA (LPC: 16:0; 18:0; 18:1; and 20:4). **(B)** Experimental design. Islets were isolated and incubated overnight in RPMI 1640 with 10% FBS and 7 mM glucose. Islets were switched to 5.5 mM glucose RPMI medium for 23-24 hrs. Islets (100 per mouse) were simulated with Vehicle (DMSO), 1.5 μM, or 5.0 μM LPC for 1 hr or 24 hrs prior to GSIS. Starvation period consisted of incubating islets in 1hr KRB buffer with low [3.3 mM] glucose at 37 Celsius. 15 size-matched islets per group were hand-picked for GSIS assays. **(C)** Insulin levels during GSIS using low [3.3 mM] or high [16.7 mM] glucose after 1-hr stimulation of isolated islets with Vehicle (DMSO), 1.5 μM LPC, and 5.0 μM LPC (n=3 per group). **(D)** Insulin levels during GSIS using low [3.3 mM] or high [16.7 mM] glucose after 24-hr stimulation of isolated islets with Vehicle (DMSO), 1.5 μM LPC, and 5.0 μM LPC (n=5 per group). Data represent mean +/- SEM, ns (no significance), *p≤0.05, **p≤0.01, ***p≤0.005, ****p≤0.0005. Data analyzed using 2-way ANOVA with correction for multiple comparisons using Holm-Šídák’s method.

### LPC Stimulation Disrupts β-cell Function Through Impaired GSIS

To determine the direct effects of LPC on β-cell function, we stimulated islets isolated from wildtype C57BL/6 mice with LPC and subsequently quantitated GSIS (**Fig. 6B**). The composition of the LPC used for these *in vitro* experiments was 60-65% palmitic (16:0), 25-27% stearic (18:0), and 6-8% oleic (18:1) acids, which matched the top 3 most abundant molecular species measured in isolated islets (**Fig. 6A**).

Stimulation experiments consisted of two different time points, acute (1 hr) and prolonged (24 hrs), and four different doses: 0μM (vehicle), 1.5 μM, 5 μM, and 50 μM LPC. The lower end of these doses used corresponds to the physiological plasma molarity seen healthy humans and patients with T2D [2-10 μM; or 100-300 μM Pi/L] [61]. All stimulations were carried out in low glucose concentrations [5.5 mM].

After 24 hr of 50 μM LPC stimulation, islets were visibly degranulated in culture (data not shown), likely from cellular apoptosis, and could not be used for GSIS assays. Acute (1-hr) LPC treatment impaired islet GSIS at the higher [5 μM], but not the lower [1.5 μM] dose (**Fig. 6C**). Prolonged (24 hr) stimulation with LPC impaired GSIS at both doses [1.5 μM; 5 μM] and disrupted basal insulin secretion at the higher [5 μM] dose (**Fig. 6D**). These data indicate that LPC stimulation directly induces β-cell dysfunction in isolated rodent islets at similar or slightly lower concentrations seen in the plasma of healthy individuals and patients with T2D.

## DISCUSSION

Although insulin resistance lies at the core of T2D, the rate of T2D progression is driven by the gradual demise of pancreatic β-cell function and/or mass. By the time hyperglycemia becomes clinically apparent, ∼80% of β-cell function will have already been lost [62]. Thus, a major goal of T2D research is to identify the mechanisms underlying β-cell dysfunction that will facilitate the design of novel treatment strategies to preserve long-term β-cell health.

In this study, we show that a 5-week course of the SGLT2i canagliflozin – the latest class of anti-hyperglycemic medications in the T2D repertoire – leads to improvements in body weight, fasting glucose, and glucose tolerance in DIO mice. These results are largely consistent with data from studies conducted in humans with impaired fasting glucose (IFG) or T2D, which demonstrate clear benefits of SGLT2i therapy on plasma glucose concentrations [33, 63, 64] and muscle insulin sensitivity [65]. By using established indices of insulin secretory capacity (e.g., Insulin 0-30 min AUC / Glucose 0-30 min. AUC; and insulin secretion / insulin sensitivity, i.e., Disposition Index), we also show that *in vivo* β-cell function is improved in DIO mice treated with SGLT2i. Several human interventional studies have demonstrated significant improvements in β-cell glucose sensitivity and secretory function *in vivo*, as measured by the incremental AUC of plasma C-peptide concentrations during hyperglycemic clamps (the gold-standard assessment of β-cell function *in vivo*) and the Disposition Index [33, 63, 64]. However, this is not a universal finding, and a very recent study in humans with pre-diabetes failed to show improvements in insulin secretion following 8-weeks of SGLT2i treatment [66].

A central question is whether improvements in insulin secretion *in vivo* following the normalization of glycemia with SGLT2i treatment reflect a durable improvement in the function of individual β-cells. In the current study, we demonstrate that SGLT2i treatment has no beneficial effect on isolated β-cell function, assessed using gold-standard GSIS assays. There are several possible explanations for the discrepancy between our *in vivo* and *in vitro* data. The effects of SGLT2i on insulin secretion *in vivo* in humans and rodents are typically assessed in the context of modest weight loss and improvements in hepatic and peripheral insulin sensitivity, a metabolic milieu that is not recreated in GSIS experiments carried out *in vitro*. Given that we and others have been unable to detect SGLT2 expression in β-cells as was originally reported in 2015 [41, 42, 67, 68], the effects of SGLT2i on insulin secretion *in vivo* are almost certainly indirect and may reflect compensatory responses that do not necessarily involve intrinsic improvements in the secretory function of individual β-cells. For example, it’s possible that SGLT2i support islet function *in vivo* by promoting α-to-β-cell transdifferentiation through these indirect mechanisms, as has recently been demonstrated in diabetic rodents [69].

The few studies that have examined GSIS in isolated β-cells following SGLT2i treatment have reported contrasting results, with some studies demonstrating an improvement [35, 70-72] and others demonstrating no benefit [73]. These discrepant findings may reflect differences in experimental design, including the dose and duration of SGLT2i therapy. However, we used a longer course of SGLT2i administration (5-weeks) compared to a prior study (4-weeks) that reported positive effects of SGLT2i on isolated β-cell function [71]. The age and the disease stage at which SGLT2i therapy is initiated could also be an important factor. One study found that a short, 2-week course of SGLT2i in db/db mice led to greater improvements in isolated β-cell function when SGLT2i therapy was started early (7 weeks of age) versus later in life (9 weeks of age) [70]. We initiated SGLT2i treatment at 16 weeks of age following an 8-week HFD, which may reflect a more advanced stage of β-cell failure in T2D. It would be of interest to systematically test whether early intervention with SGLT2i reverses β-cell dysfunction in models of T2D. Consistent with the findings of the present study, Jurczak and colleagues demonstrated that deletion of SGLT2 in db/db mice improved glucose homeostasis and indices of β-cell function *in vivo*, but failed to enhance GSIS in isolated islets [74]. The authors concluded that increased islet mass (60% higher in SGLT2KO vs control), not individual islet function *per se*, accounted for the improvement in glucose homeostasis and β-cell function observed *in vivo* [74]. Our islet size distribution analysis demonstrated a ∼40% increase in the average size of islets isolated from HFD-fed mice. However, islet area was restored following SGLT2i treatment in our study, suggesting that, while it is feasible that SGLT2i treatment influences β-cell mass *in vivo*, alternative mechanisms likely underlie persistent β-cell dysfunction following the normalization of glycemia in these animals.

A significant finding of our study is that islet lipotoxicity contributes, at least in part, to impaired β-cell function in rodent models of T2D. Most of the lipidome changes we observed in HFD-fed mouse islets occurred in glycerophospholipids in addition to arachidonic acid (20:4) species. Glycerophospholipids are the major structural lipid component of eukaryotic cell membranes and serve an important function in cell growth, migration, and signal transduction. Phosphatidylcholine (PC) and PE account for more than 50% of glycerophospholipids, and alterations in PC and PE levels and/or their ratio has consistently been linked to metabolic abnormalities, including insulin resistance in skeletal muscle [52] and adipose tissue [75]. We observed elevated LPC, which is derived from PC in lipoproteins and/or through cleavage of PC via PLA2 [57, 76], in islets isolated from HFD-fed mice before and after SGLT2i. To confirm a mechanistic impact of LPC on β-cell function, we treated healthy isolated islets with LPC. These experiments clearly demonstrated that LPC negatively impacts GSIS, which is consistent with our lipidomics data [77]. LPC stimulation has been previously shown to enhance GSIS in several insulinoma cell lines [78, 79], but not in primary rodent islets. While our data showed an increase in basal insulin secretion with prolonged exposure to LPC, GSIS was inhibited in a dose- and time-dependent manner. This is consistent with the cytotoxic effects of LPC, which induces inflammatory damage in endothelial cells [80], disrupts mitochondrial function in primary hepatocytes [81], and triggers the release of pro-inflammatory cytokines from adipocytes [82] and immune cells [83]. Additional studies will be needed to further evaluate the pathophysiological role of LPC in β-cell failure in T2D.

Following HFD-feeding, we observed a significant increase in some lipid species long considered lipotoxic in T2D (i.e., ceramides), but not others (i.e., TAGs and DAGs). In the islets of Zucker diabetic fatty (ZDF) rat model of T2D, Unger and colleagues demonstrated that islet ceramide content was significantly elevated and linked to β-cell apoptosis [84]. Recent studies using alternative rodent models of T2D have shown only modest or no increases in β-cell ceramide content *in vivo* [85, 86]. Our data support the findings from Unger et al. and suggest that ceramides have an important lipotoxic role in DIO mice. The role of TAG accumulation in β-cell failure in T2D remains controversial. The importance of pancreatic TAG in the pathogenesis of T2D was initially demonstrated in rodents [87]. In human patients with T2D, weight loss secondary to gastric bypass surgery reduced excess pancreatic TAG and improved first phase insulin secretion in response to a stepped glucose infusion [88]. However, these data reflect total pancreatic TAG assessed using MR spectroscopy and, to the best of our knowledge, no other study has examined TAG concentrations specifically in islets isolated from humans or rodents. In future studies, we hope that the shotgun lipidomics methodology employed herein will facilitate the quantification of lipid species in islets isolated from animal and human models of T2D.

An important caveat of the experimental model employed in the current study is illustrated in a recent article published by Marselli et al. [89]. In this study, the authors demonstrate the differential effects of distinct metabolic stressors on β-cell function and indicate that some unique combinations of glucolipotoxic stimuli are reversible, whereas others persist even after the stressor is removed. In this context, it is possible that, in our experiment, hyperglycemia during the HFD irreversibly damaged β-cells, which led to persistent β-cell dysfunction even after the removal of this stressor using SGLT2i. Put simply, our SGLT2i intervention was “too little, too late”. However, the same argument could be made for lipotoxicity during the HFD, and it is possible that lipid species partially or completely normalized following SGLT2i treatment (i.e., ceramides) irreversibly damaged β-cell function during the HFD period. Despite our data demonstrating a direct effect of LPC on isolated β-cells, it is plausible that multiple lipid species contribute to the lipotoxic signal during overnutrition in both humans and rodents, but careful follow-up studies will be needed to tease out the roles of individual lipids *in vivo*.

Taken together, our study provides novel insights into the role of lipotoxicity in the pathogenesis and treatment of β-cell failure in T2D. We have identified a pro-inflammatory lipid, LPC, as a potential culprit for sustained β-cell lipotoxicity in the absence of glucotoxicity. If these findings can be replicated in human models, they suggest that early interventions targeting lipotoxicity may be necessary to protect and/or restore β-cell function in patients with T2D.

## EXPERIMENTAL PROCEDURES

### Animal Husbandry and Experimental Design

Eight-week-old wild-type (C57BL/6J; JAX #000664) male mice were purchased from The Jackson Laboratory, housed in environmentally-controlled conditions (23°C, 12-hour light/dark cycles) and provided ad-libitum access to water and a regular chow (RC; 70% kcal carbohydrate, 10% kcal fat, 20% kcal protein; D12450J; Research Diets Inc.) or high-fat diet (HFD; 60% kcal from fat, Research Diets, D12492) for 8 weeks. Mice were subsequently divided into the following diets and treatment groups for 5 weeks: (a) RC + vehicle (cellulose, 10 mg/kg/day); (b) HFD + vehicle; (c) HFD + CANA (canagliflozin; 10 mg/kg/day). Administration of vehicle or CANA was administered daily via oral gavage. Food consumption and body weight were monitored weekly for 13 weeks, and blood glucose levels were measured daily during the 5-week therapy just prior to gavage. All procedures were approved by the Institutional Animal Care and Use Committee (IACUC) at University of Texas Health San Antonio (UTHSA).

### Glucosuria measurements

Immediately after oral gavage of vehicle or CANA, mice were placed in metabolic cages with ad libitum water and no food for 5 hours. Urine was collected afterwards, and urine glucose was measured using Amplex™ Red Glucose/Glucose Oxidase Assay kit (Invitrogen; catalog #: A22189).

### Pancreatic Intraductal Injections and Mouse Islet Isolation

Islets were isolated as described previously [90] using the pancreatic intraductal collagenase technique with slight modifications. Briefly, the common bile duct was cannulated in the anterograde direction and the pancreas distended with CIzyme RI (Vitacyte; catalog #: 005-1030). Pancreases were harvested and digested for 17 min at 37°C. Islets were isolated by generating sedimentation gradient of Lymphocyte Separation Medium (Corning; catalog #: 25-072-CV). Floating islets were washed, and healthy islets were hand-picked and transferred into RPMI 1640 (Thermo Fisher) islet culture medium containing 7 mmol/l glucose, 10% fetal bovine serum (FBS), and 1% penicillin-streptomycin at 37°C in 5% CO_2_ for overnight recovery incubation.

### Glucose Stimulated Insulin Secretion (GSIS) Assays

After 20-24hrs of incubation in RPMI 1640 with 7mM glucose, islets were washed with low (3.3 mM) glucose KRB medium and starved by culturing them for 1 hr. at 37°C using the same low glucose medium. For secretion experiments, 10 size-match islets (120–150 μm) were handpicked from each mouse and transferred into a 1.5mL Eppendorf tube. Islets were exposed to KRB medium containing low (3.3 mM) followed by high (16.7 mM) glucose. Next, islets were depolarized with KRB containing 30mM KCl. Low (3.3 mM) glucose KRB medium was used to wash islets in between all steps. Finally, islets were sonicated, and total insulin content in the supernatant was collected for normalization. Insulin concentrations were measured by ELISA (Ultra-Sensitive Mouse Insulin ELISA kit, Chrystal Chem, Downers Grove, IL, USA; catalog #: 90080) following manufacturer’s instructions.

### Quantification of Pancreatic Islet Size Distribution

Optical images of islets in culture were obtained with a IX2-SLP2 Olympus brightfield microscope using 10X objective lens. Islet size analysis was performed using ImageJ software. Between 250-350 islets per group were manually circled and quantified. A hemocytometer was used for reference to set the scale in the microscope at 12.68 pixels / μM.

### Intraperitoneal Glucose Tolerance Test (IPGTT) and Serum Insulin Quantifications

Mice were fasted for 5-6 hrs. and IPGTTs were performed. Blood samples were drawn from the tail vein before injecting a fixed dose (50 µg in 500 µl) of a 10% dextrose solution (B. Braun; catalog #: L5202; t = 0 min) and after 15, 30, 60, and 120 minutes. Glucose levels were measured using an automated glucose meter. Serum insulin was collected during IPGTT to assess insulin response. Insulin quantification was performed with Luminex Kit (Millipore, MILLIPLEX MAP Mouse Metabolic Hormone Magnetic Bead Panel - Metabolism Multiplex Assay; catalog #: MMHMAG-44K) following manufacturer’s instructions.

### Immunohistochemistry (IHC)

Pancreases were harvested, fixed with Formalin (4% paraformaldehyde) over night at 4°C, embedded in paraffin, and sectioned at 5μm. Slides were deparaffinized and rehydrated. For antigen retrieval, tissue sections were then heated in a microwave (maximum power for 1 min., and 20% power for 10 min.) in citrate buffer (pH 6.0), followed by permeabilization and blocking with 5% donkey serum in PBS with or without 0.1% Triton X-100 for 1 hour at room temperature (r.t.). Primary antibodies were diluted in antibody diluent (Dako) and incubated overnight at 4°C. Secondary antibodies were diluted in PBS and incubated for 1 hour at r.t. DAPI was used to stain cell nuclei. Slides were mounted with Cytoseal (Thermo) and images were captured on an LAS X confocal microscope. Antibodies used are provided in **Supplemental Table 2**. Quantification analyses were carried out manually.

### Islet Lipidomics Methodology

Islets were lyophilized and homogenized in ice-cold diluted phosphate-buffered saline (0.1X PBS). Lipids were extracted by a modified procedure of Bligh and Dyer extraction in the presence of internal standards, which were added based on total protein content as previously described [91, 92]. A triple-quadrupole mass spectrometer (Thermo Scientific TSQ Altis, CA, USA) and a Quadrupole-OrbitrapTM mass spectrometer (Thermo Q ExactiveTM, San Jose, CA) equipped with a Nanomate device (Advion Bioscience Ltd., NY, USA) and Xcalibur system software was used as previously described [93]. Briefly, diluted lipid extracts were directly infused into the electrospray ionization source through a Nanomate device. Signals were averaged over a 1-min period in the profile mode for each full scan MS spectrum. For tandem MS, a collision gas pressure was set at 1.0 mTorr, but the collision energy varied with the classes of lipids. Similarly, a 2- to 5-min period of signal averaging in the profile mode was employed for each tandem MS mass spectrum. All full and tandem MS mass spectra were automatically acquired using a customized sequence subroutine operated under Xcalibur software. Data processing including ion peak selection, baseline correction, data transfer, peak intensity comparison, 13C deisotoping, and quantitation were conducted using a custom programmed Microsoft Excel macro as previously described after considering the principles of lipidomics [94]. Lipid amount data was normalized to islet number given significant differences in islet size and total protein content between HFD and RC islets.

### *In Vitro* LPC Stimulations

Pancreatic islets from wild-type, C57BL6 mice at 25-30 weeks of age were isolated as described above and maintained in RPMI 1640 medium supplemented with 10% FBS overnight. LPC stimulations were carried out in this medium using 100 islets per mouse. Islets were stimulated for 1hr (acute) or 24 hours (long-term) at 0.0μM (Vehicle DMSO), 1.5μM, 5.0μM or 50μM concentrations at 37° C. After LPC stimulation, 15 size-match islets (120–150 μm) were manually picked for GSIS assay as described above.

### Statistical Analyses

Data are expressed as mean ± standard error of the mean (SEM). To confirm significance, Welch’s t test (accounting for unequal sample sizes and variances) was used for comparisons of 2 groups. Data containing three or more groups was analyzed using 1-way or 2-way ANOVA (adjusted for repeat measures, where appropriate) with Holm-Sidak post-hoc tests accounting for multiple comparisons, where appropriate. Significance was set at p ≤ 0.05. Statistical analyses were performed using PRISM 9 (GraphPad Software).

### Study Approval

All animal experiments were conducted after approval by the Institutional Animal Care and Use Committee (IACUC) of The University of Texas Health San Antonio in accordance with National Institutes of Health (NIH) guidelines.

## Data and Resource Availability

Data sets generated during the current study are available from the corresponding author upon reasonable request.

## AUTHOR CONTRIBUTIONS

Conceptualization, I.A.V., and L.N.; Methodology, I.A.V., J.P.P., and L.N.; Formal analysis, I.A.V., J.P.P. and L.N.; Investigation, I.A.V., A.K., T.M.B., C.E.S., M.F., I.A., and Z.X.; Writing – Original Draft, I.A.V., and L.N.; Writing – Review and Editing, I.A.V., C.S., X.H., and L.N.; Supervision, L.N.; Funding Acquisition, L.N.

## ACKNOWLEDGEMENTS

This work was supported by the American Diabetes Association (grant #: 1-19-PMF-007) and the National Institutes of Health (grant #: R01DK128247)

