## Supplementary Figures for "Persistent Inflammatory Lipotoxicity Impedes Pancreatic β-cell Function in Diet-Induced Obese Mice Despite Correction of Glucotoxicity"

**SUPPLEMENTAL INFORMATION**

### SUPPLEMENTARY FIGURE LEGENDS

**Supp. Figure 1.**

**(A)** Total AUC for 0-to-30-minute insulin levels (I30) during post-treatment IPGTT **(Fig. 2G)** (n=7-8 per group).

**(B)** Total AUC for 0-to-30-minute glucose levels (G30) during post-treatment IPGTT **(Fig. 2D)** (n=7-8 per group).

Data represent mean +/- SEM; ns (no significance), *p≤0.05, **p≤0.01, ***p≤0.005, ****p≤0.0005. Data analyzed using an ordinary 1-way ANOVA with correction for multiple comparisons using Holm-Šídák's method.

**Supp. Figure 2.**

**(A)** Representative images of immunohistochemistry experiments on paraffin-embedded pancreatic sections of RC and HFD mice; SGLT1 or SGLT2 (Red), Insulin (green); DAPI (blue).

**(B)** Representative images of immunohistochemistry experiments on isolated wild-type islets; SGLT21 or SGLT2 (Red), Insulin (green); DAPI (blue).

**Supp. Figure 3.**

**(A)** Total islet insulin content in 10 islets used for GSIS (**Fig. 3F**) (n=10-12 per group).

**(B)** Insulin levels during GSIS after stimulation of isolated islets with 30mM KCL (n=5 per group).

**(C)** Insulin levels during GSIS using low [3.3mM] or high [16.7mM] glucose (n=5-7 per group).

Data represent mean +/- SEM; ns (no significance). *p≤0.05, **p≤0.01, ***p≤0.005, ****p≤0.0005. Data analyzed using Brown-Forsythe 1-way ANOVA, with correction for multiple comparisons using Dunnett's method **(Supp. Fig. 3A)** and 2-way ANOVA with correction for multiple comparisons using Turkey's method **(Supp. Fig 3B-C)**.

**Supp. Figure 4.**

**(A)** Representative images of immunohistochemistry (IHC) experiments on pancreatic sections using secondary antibodies alone; Alexa-594 [1:100] (Red), Alexa-488 [1:100] (green); DAPI (blue). Scale = 100μM.

**Supp. Figure 5.**

**(A**) Individual molecular species for Fatty Acyl Lipids (FFAs and Acyl-CARs).

**(B)** Individual molecular species for Glycerolipids (DAGs and TAGs). Note: Only the top 20 (of 87) most abundant molecular species detected for TAG are shown.

**(C)** Individual molecular species for Sphingolipids (SMs and CERs).

**(D)** Individual molecular species for Glycerophospholipids (PE, LPE, PC, CL, PG, PG, PI, PS, PA). Note: Data for LPC shown in Fig. 5A. Data for PI consisted of 1 molecular species (PI 18:0-20:4), which is shown in **Fig. 5F** as total PI.

**(D')** Absolute amount of PE broken down into Diacyls and Plasmalogens.

**(E-F)** Heatmap depicting fold change (z-scores) for significantly elevated glycerophospholipid **(E)** and non-glycerophospholipids **(F)** species in isolated islets from RC vs HFD mice. Lipids in the upper box were elevated in HFD and remained elevated after CANA treatment. Lipids in the lower box were elevated in HFD but normalized by CANA treatment.

Data represent mean +/- SEM; ns (no significance), *p≤0.05, **p≤0.01, ***p≤0.005, ****p≤0.0005. (n=4-6 per group) Data analyzed using an ordinary 2-way ANOVA with correction for multiple comparisons using Holm-Šídák's method. All data normalized to islet number.

**Supp. Figure 6.**

**(A**) Relative abundance of Fatty Acyl Lipids (FFAs and Acyl-CARs).

**(B)** Relative abundance of Glycerolipids (DAGs and TAGs).

**(C)** Relative abundance of Sphingolipids (SMs and CERs).

**(D)** Relative abundance of Glycerophospholipids and Lyso-glycerophospholipid intermediates (LPE, PC, PS, PI, PG, PA, CL, Lyso-CL). Note: Relative abundance data for PE and LPC are shown in **Fig. 5L and 5M**, respectively.

Data represent mean +/- SEM; ns (no significance). *p≤0.05, **p≤0.01, ***p≤0.005, ****p≤0.0005. (n=4-6 for all groups). Data analyzed using Brown-Forsythe 1-way ANOVA, with correction for multiple comparisons using Dunnett's method. All data normalized to islet number.

Supp. Figure 1

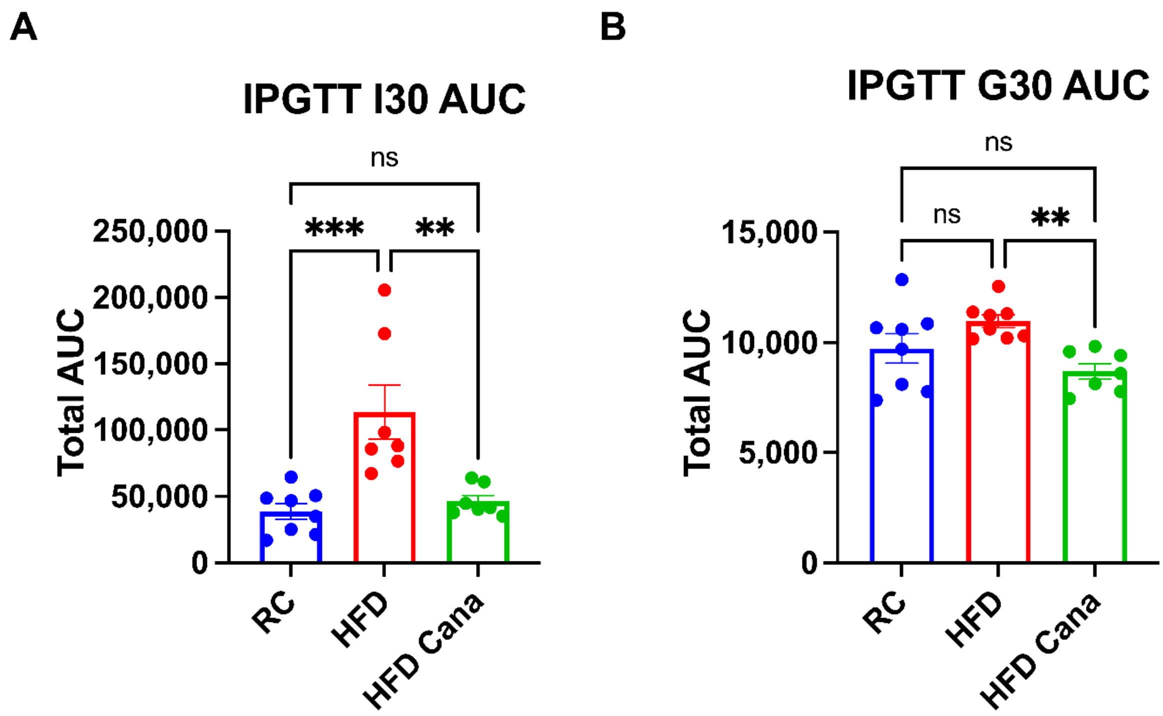

Supp. Figure 2

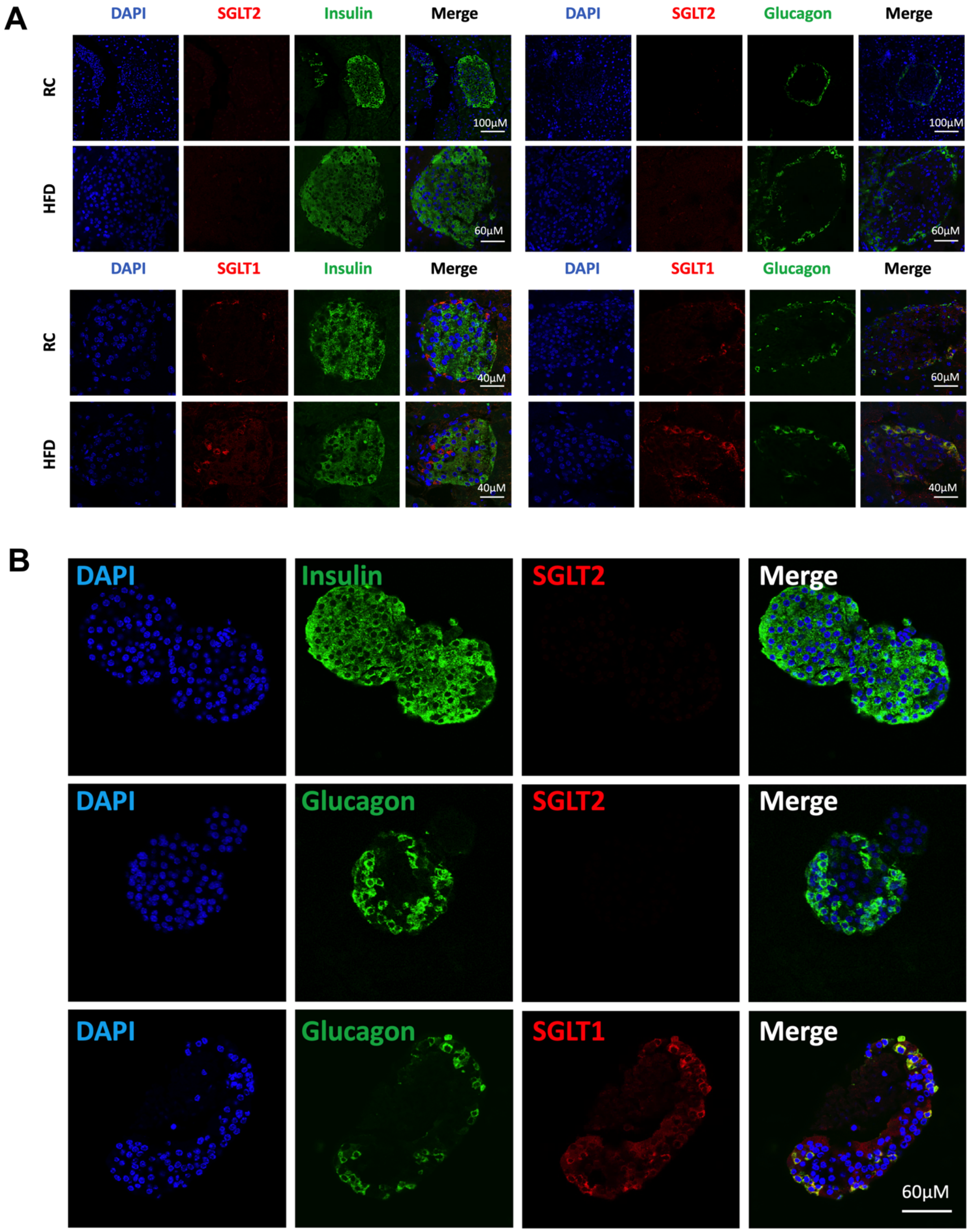

Supp. Figure 3

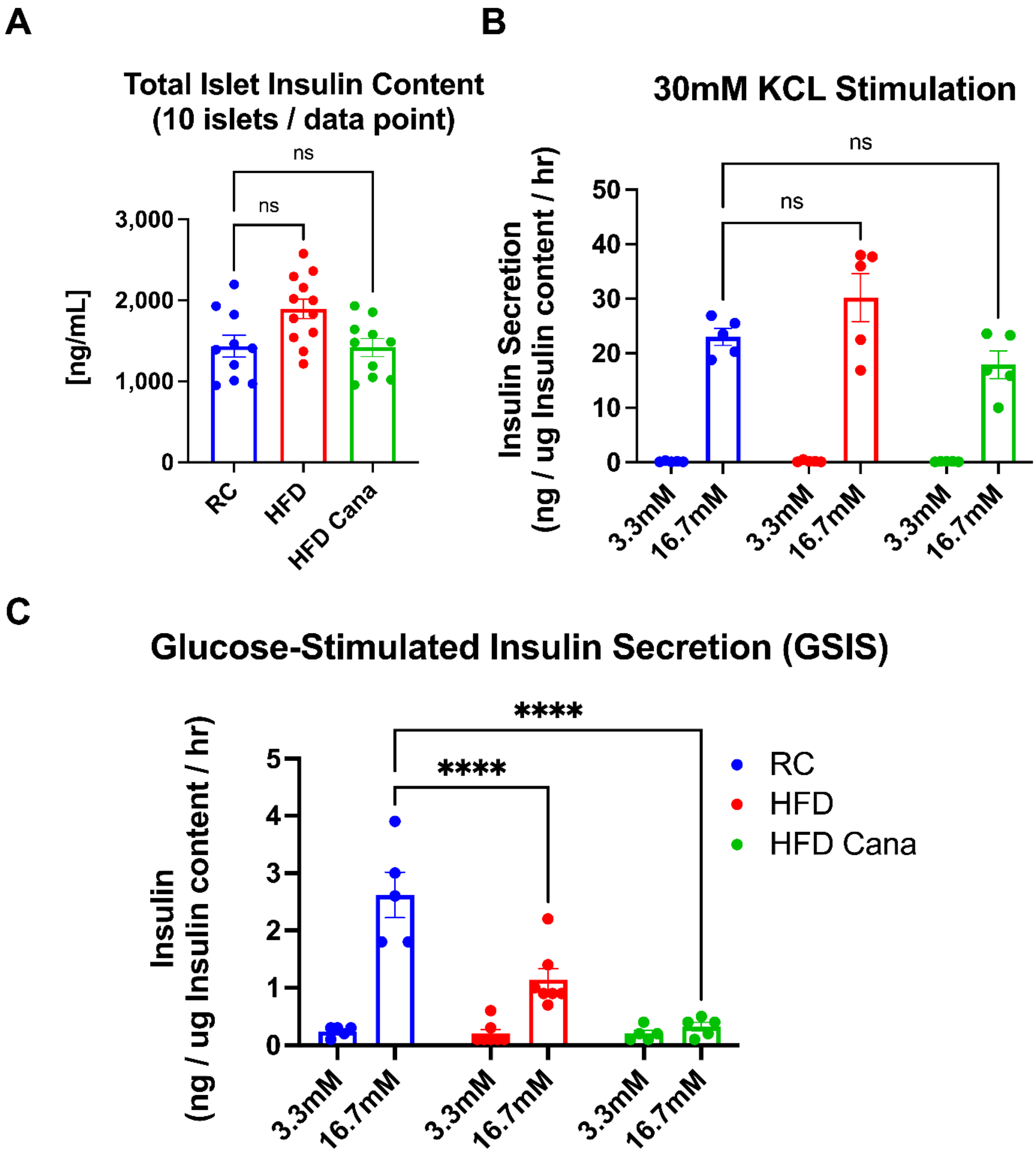

**Supp. Figure 4**

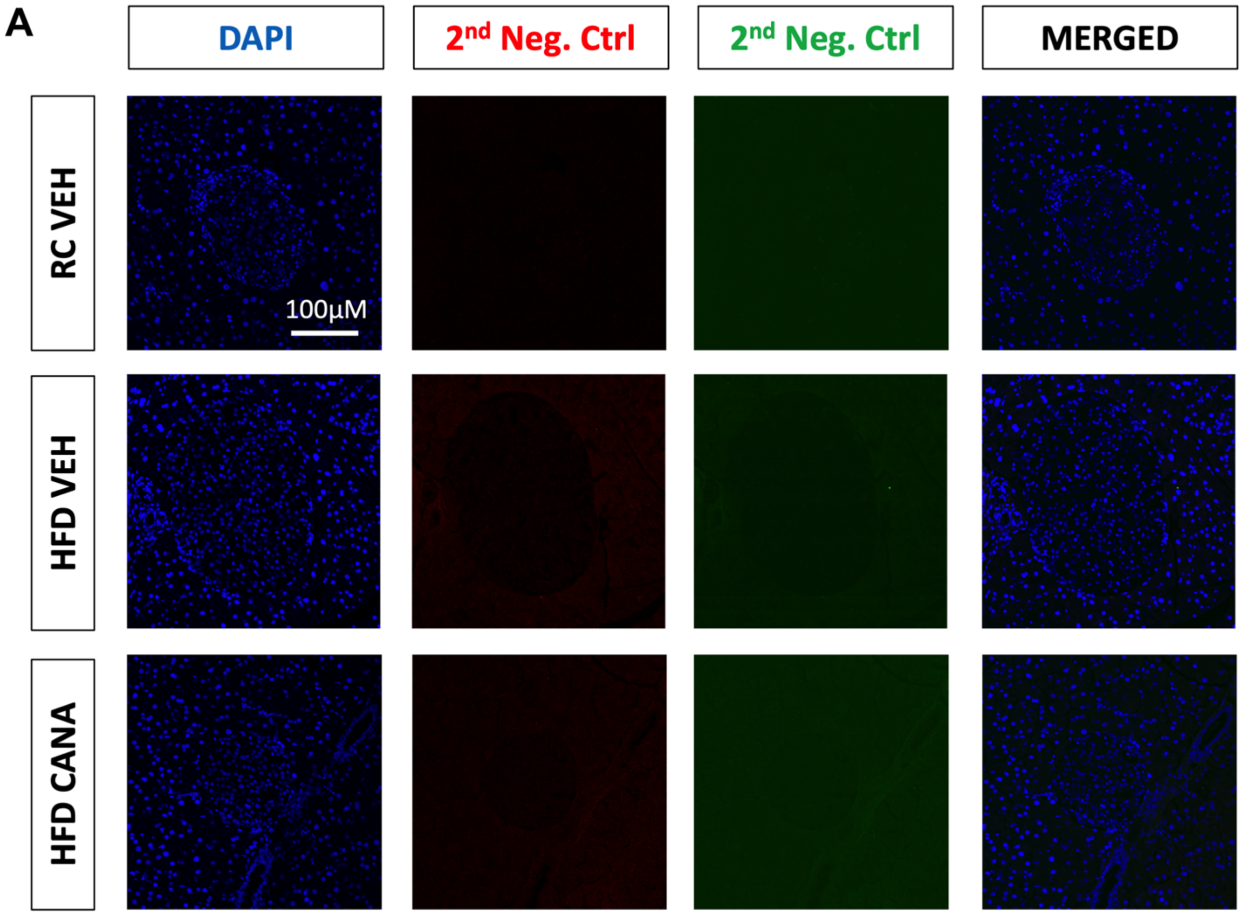

**Supp. Figure 5**

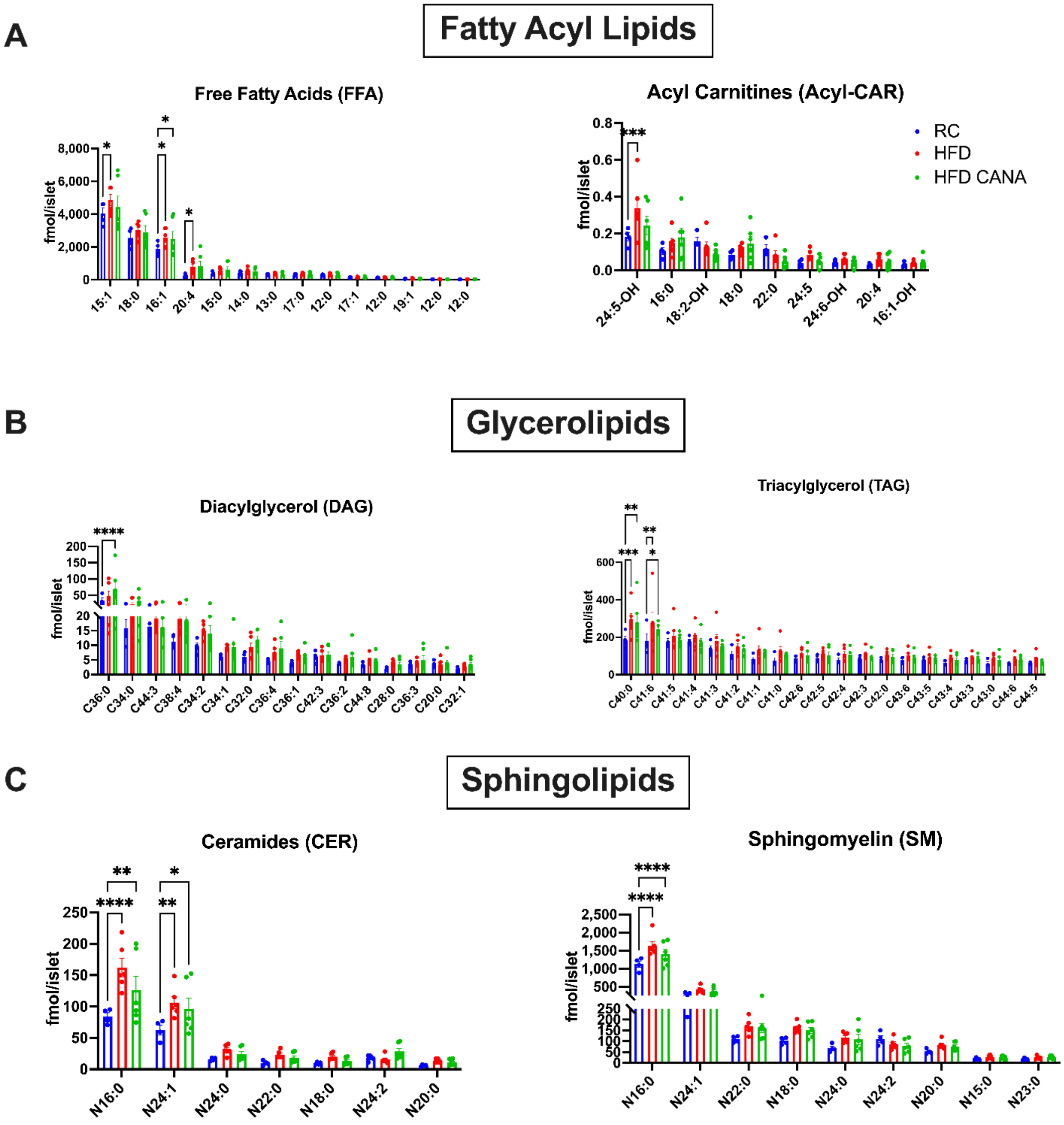

**Supp. Figure 5 (Cont'd)**

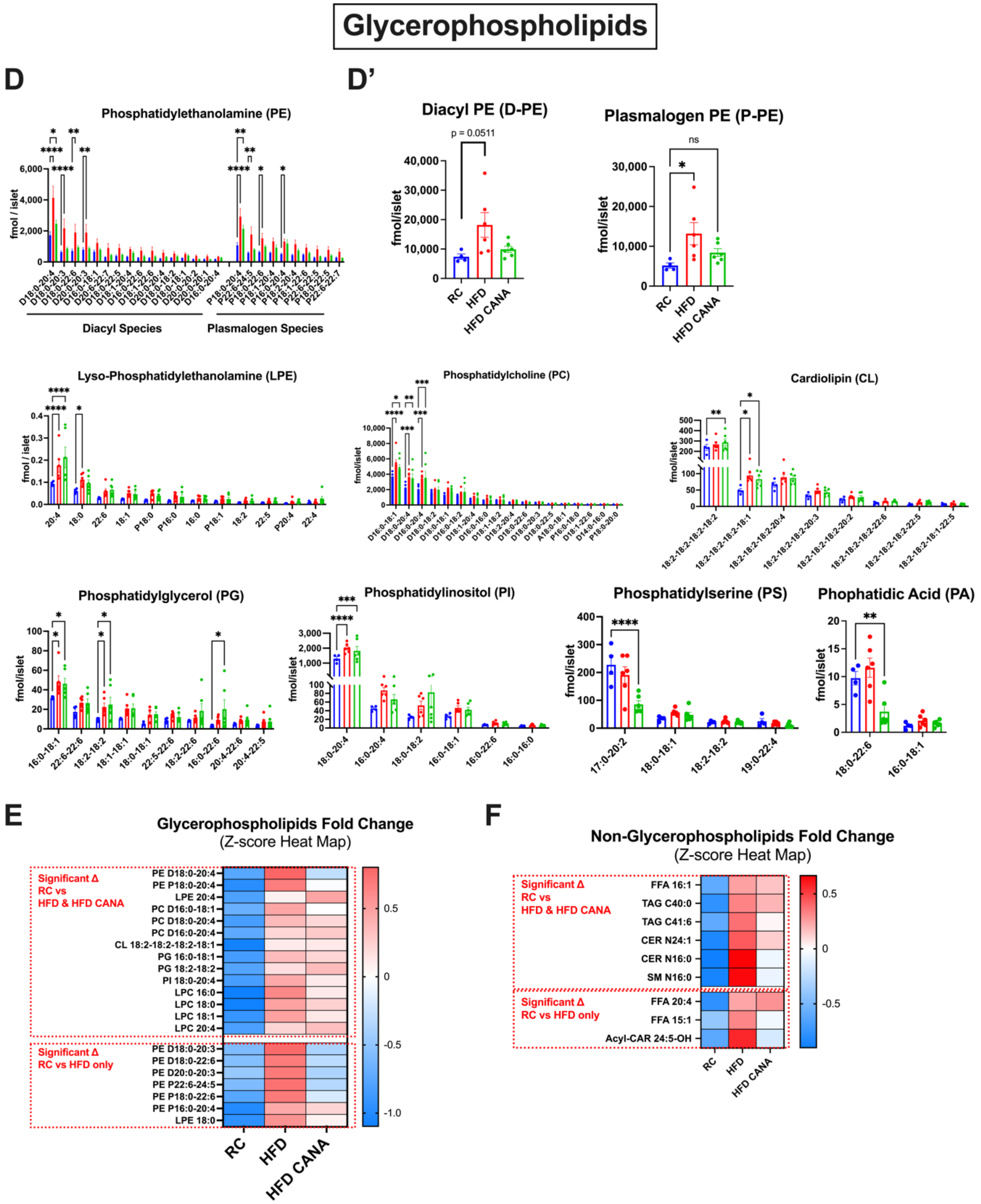

Supp. Figure 6

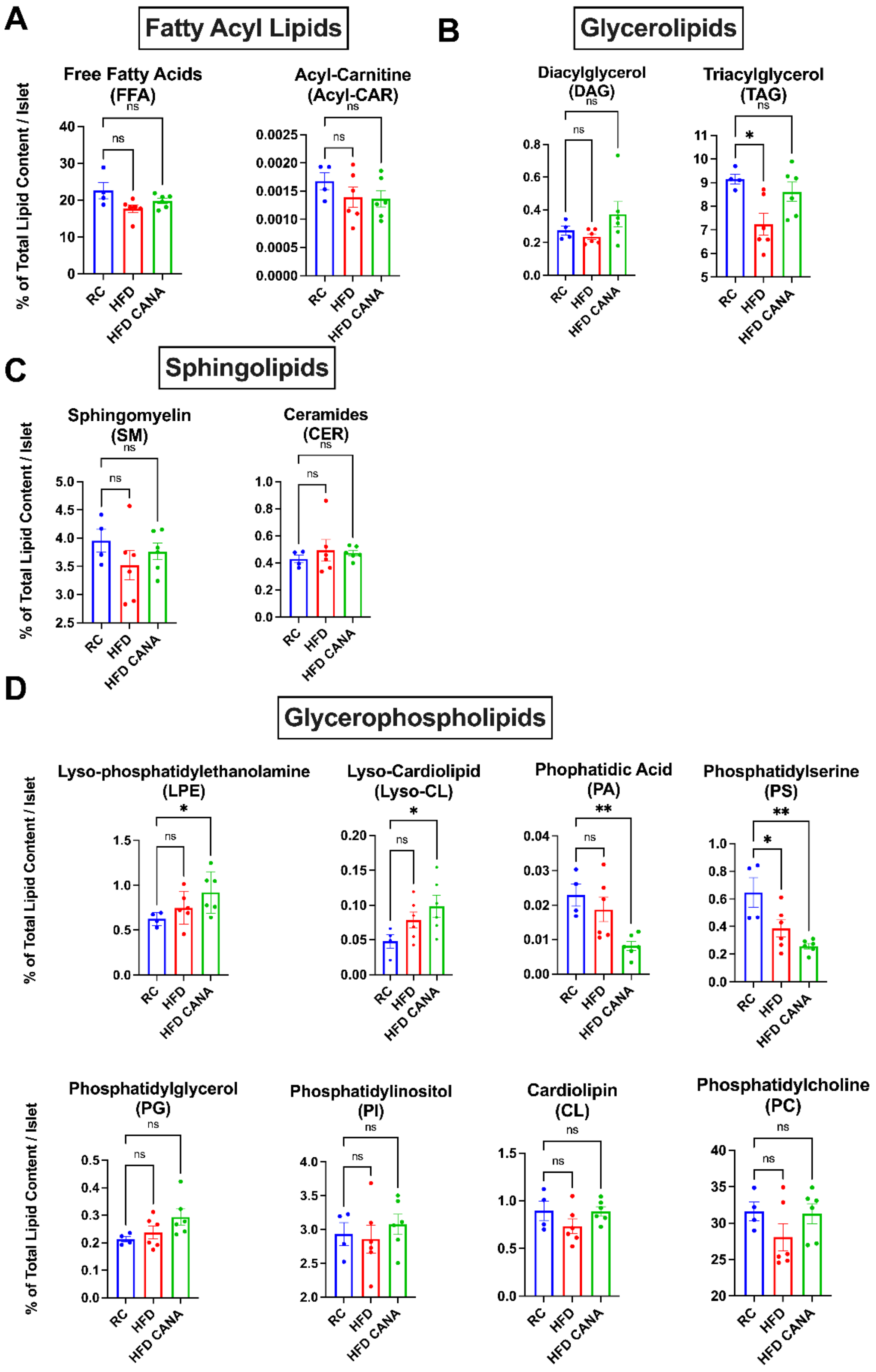

**Supp. Table 1.** Samples used for lipidomics analysis.

| | Sample Number | Islet Number |
|---|---|---|
| RC Vehicle (n=4) | Sample 1 | 410* |
| | Sample 2 | 360* |
| | Sample 3 | 300* |
| | Sample 4 | 340* |
| HFD Vehicle (n=6) | Sample 1 | 320* |
| | Sample 2 | 320* |
| | Sample 3 | 370 |
| | Sample 4 | 360 |
| | Sample 5 | 300 |
| | Sample 6 | 310* |
| HFD Cana (n=6) | Sample 1 | 310* |
| | Sample 2 | 310 |
| | Sample 3 | 380 |
| | Sample 4 | 300 |
| | Sample 5 | 320 |
| | Sample 6 | 300* |

* = sample is an aggregate of 2 mice

**Supp. Table 2.** Antibodies in the present study.

| **Species** | **Primary Antibody** | **Catalog Number** | **Company** | **Concentration** |
|---|---|---|---|---|
| Guinea Pig | Insulin | Ab7842 | Abcam | [1:100] |
| Mouse | Glucagon | SC-57171 | Santa Cruz | [1:100] |
| Goat | SGLT1 | Ab240474 | Abcam | [1:100] |
| Rabbit | SGLT1 | 07-1417 | EMD Millipore | [1:100] |
| Rabbit | SGLT2 | 14210S | Santa Cruz | [1:100] |
| Rabbit | SGLT2 | Ab37296 | Abcam | [1:100] |
| Rabbit | CD68 | 76437S | Santa Cruz | [1:100] |

| **Secondary Antibodies** |
|---|
| Alexa Fluor® 488 Goat anti-pig IgG (A11073, Life Technologies) 1:400 |
| Alexa Fluor® 488 Chicken anti-mouse IgG (A21200, Life Technologies) 1:400 |
| Alexa Fluor® 488 Goat anti-rabbit IgG (A11008, Life Technologies) 1:400 |
| Alexa Fluor® 568 Donkey anti-rabbit IgG (A11008, Life Technologies) 1:400 |
